# The Genome of the American Dog Tick (*Dermacentor variabilis*)

**DOI:** 10.1101/2025.03.12.642860

**Authors:** Jacob Cassens, Matt Villalta, Saul Aguirre, Lauren Ecklund, Trek Stenger, Idil Abdi, Sree Venigalla, Elizabeth Shiffman, Kristen Bastug, Beth K. Thielen, Christopher Faulk

**Author notes:** Denotes corresponding author. Contact. **Contacts** Jacob Cassens Matt Villalta Saul Aguirre Lauren Ecklund Trek Stenger Idil Abdi Sree Venigalla Elizabeth Shiffman Kristen Bastug Beth K. Thielen Christopher Faulk.

## Abstract

The American dog tick (*Dermacentor variabilis*) is a vector of zoonotic pathogens in North America that poses emerging threats to public health. Despite its medical importance, genomic resources for *D. variabilis* remain scarce. Leveraging long-read nanopore sequencing, we generated a high-quality genome assembly for *D. variabilis* with a final size of 2.15 Gb, an N50 of 445 kb, and a BUSCO completeness score of 95.2%. Comparative BUSCO analyses revealed fewer duplicate genes in our assembly than in other *Dermacentor* genomes, indicating improved haplotype resolution. The mitochondrial genome, assembled as a single circular contig, clustered monophyletically with *D. variabilis* isolates from the Upper Midwest, corroborating regional phylogenetic relationships. Repetitive element analysis identified 61% of the genome as repetitive, dominated by LINEs and LTR elements, with 24% remaining unclassified, underscoring the need for further exploration of transposable elements in tick genomes. Gene annotation predicted 21,722 putative genes, achieving a protein BUSCO completeness of 80.88%. Additionally, genome-wide methylation analysis revealed 9.9% global 5mC methylation, providing the first insights into epigenetic modifications in *D. variabilis*. Further, nanopore sequencing detected *Rickettsia montanensis* and a non-pathogenic *Francisella*-like endosymbiont. These findings expand our understanding of tick genomics and epigenetics, offering valuable resources for comparative studies and evolutionary analyses.

## Introduction

*Dermacentor variabilis* (Say, 1821), also known as the American dog tick or wood tick, is an obligate blood-feeding ectoparasite found throughout North America, first described over 200 years ago (Lado et al., 2019). The American dog tick has three post-embryonic life stages (larva, nymph, adult), each requiring a blood meal for development and reproduction. Each life stage provides an opportunity to acquire a blood meal, and *D. variabilis* is a three-host tick that can feed on a different host species at each life stage. The immature life stages commonly parasitize smaller mammals, such as rodents, while adults primarily feed on larger animals, including dogs and deer (Duncan et al., 2021). This blood-feeding strategy facilitates the enzootic maintenance of tick-borne pathogens. The American dog tick is a vector for human pathogenic Rocky Mountain Spotted Fever (*Rickettsia rickettsii*) and Tularemia (*Francisella tularensis*). It has also been found infected with other bacterial (*Anaplasma marginale, Ehrlichia* spp.), protozoal (*Toxoplasma gondii, Cytauxzoon felis, Babesia* spp.), and viral (Powassan) pathogens, although its vector competency for these agents is unknown (Ben-Harari, 2019; Duncan et al., 2021; Hart et al., 2023; Reichard et al., 2021; Shock et al., 2014; Steiert and Gilfoy, 2002; Whitten et al., 2019).

Additionally, *D. variabilis* exhibits aggressive host-seeking behavior and is most active during times of the year with increased human outdoor activity (e.g., April-September), making it one of the most frequently encountered hard ticks in the US (Duncan et al., 2021). Although pathogen prevalence in natural populations remains low (Eisen et al., 2017), expanding genomic resources for hard ticks like *D. variabilis* is critical to generating fundamental insights into tick biology and addressing global zoonotic threats.

Ticks present unique challenges to genomic research due to their large genome sizes, high proportion of repetitive elements, and complex life cycles. Genomic studies on *D. variabilis* and other ticks have been limited yet are critical as ticks continue to expand their geographic ranges and pose growing threats to public health. The American dog tick belongs to the class Arachnida, order Ixodida, family Ixodidae, and group Metastriata. Metastriate ticks comprise 13 genera with over 450 species, representing a derived lineage relative to the basal Prostriata (Mans et al., 2016). Currently, only ten unique genome assemblies for metastriate ticks are available on NCBI, and these resources have proved valuable for investigating tick biology and evolution. For example, comparative studies of metastriate and prostriate genomes in Europe and Asia have highlighted lineage-specific evolutionary trajectories of tick-host interactions, tick immunity, and ecological adaptations (Cerqueira De Araujo et al., 2025; Jia et al., 2020).

The recent advancements in long-read sequencing technologies, such as Oxford Nanopore Technologies (ONT) and Pacific Biosciences (PacBio), have revolutionized tick genomics by overcoming challenges associated with assembling large and repetitive genomes. Our study contributes to this growing field by presenting the first whole-genome assembly for *D. variabilis*. This high-quality genome assembly provides a foundational resource for comparative genomic analyses, population genetics, and evolutionary studies. Further, as obligate blood feeders, ticks are a repository for exogenous DNA– previous blood meals, endosymbionts, and pathogens–that is recoverable from a single sample using native genomic sequencing. We demonstrate the utility of long-read sequencing technologies in differentiating tick DNA from pathogenic and non-pathogenic DNA while broadening our understanding of the genetic diversity, evolutionary history, and adaptive strategies of medically significant arthropods. Developing genomic resources for *D. variabilis* enhances the ability to identify novel targets for transgenic techniques and pest management strategies, aiding efforts to mitigate the impact of hard ticks on public health.

## Methods

### Sample collection

Adult female *D. variabilis* ticks found attached to a companion canine after an outdoor walk in central Minnesota were removed and stored in 100% ethanol. Ticks were morphologically identified using the taxonomic key from Yunker et al., 1986. Ticks were transferred to Zymo DNA/RNA shield for one hour prior to DNA extraction and library preparation.

### DNA extraction and sequencing

Genomic DNA was extracted from 10 adult female ticks using a MagAttract Blood DNA/RNA Kit (Qiagen, Venlo, Netherlands) according to the manufacturer’s instructions. Fragment size was assessed on an agarose gel. Size selection for fragments > 10 kb was attempted by treatment with a short fragment eliminator XS kit from PacBio. However, insufficient DNA was recovered for sequencing, necessitating the sequencing of short fragments only. Sequencing was performed on a P2 Solo instrument (ONT) using a single PromethION R10.4.1 flow cell. Data was collected using MinKNOW v23.07.12 in the 5 khz condition.

Raw nanopore data in pod5 format was re-basecalled using dorado v0.8.2 (https://github.com/nanoporetech/dorado) with model dna_r10.4.1_e8.2_400bps_sup@v5.0.0, and modifications called simultaneously with the flag ‘--modified-bases 5mC_5hmC’. Read quality was assessed using the Nanoq package (https://github.com/esteinig/nanoq).

### Genome Assembly

The genome was *de novo* assembled using Flye v2.9 on a high memory (2 Tb) node due to the expected assembly size and our read depth (https://github.com/fenderglass/Flye). The resulting draft assembly was processed with the NCBI Foreign Contaminant Screen program to remove vector and adapter sequences (https://github.com/ncbi/fcs). Duplicate contigs and overlaps were removed with Purge_dups v1.2.6 (https://github.com/dfguan/purge_dups). Gaps were filled in the assembly by using the ntLink v1.3.11 scaffolding program with six rounds of linking and gap-filling (https://github.com/bcgsc/ntLink). The resulting scaffolded assembly was split back into contigs at all remaining unfilled gaps, and “n” bases were removed, creating a more contiguous contig-level assembly.

The genome was decontaminated using Blobtoolkit v4.4.0 (https://blobtoolkit.genomehubs.org). First, the contigs were annotated using locally installed NCBI BLAST with the ‘core_nt’ database. Second, contig depth was calculated using samtools coverage. Finally, the blobtools database was generated and annotations analyzed in a browser. Hits assigned to “arthropoda” “no-hit” were kept and all other contigs removed. We also removed any contigs with less than 10X or greater than 1000X depth. The resulting assembly was designated as the reference.

After each assembly iteration, the completeness of draft assemblies was evaluated using the standard Benchmarking Universal Single-Copy Ortholog (BUSCO) count with the Arachnida lineage (Simão et al., 2015). We used the compleasm package (https://github.com/huangnengCSU/compleasm; Huang and Li, 2023) to generate completeness scores as it offers a faster, more accurate implementation of BUSCO. We combined the single and duplicate counts from the compleasm scores to provide a direct comparison to BUSCO’s ‘complete’ value. Genome summary statistics, such as N50 and L50, were calculated using assembly-stats (https://github.com/sanger-pathogens/assembly-stats). The quality of each draft assembly was assessed by the read N50 (i.e., contig length at which 50% of the assembled genome is contained in longer contigs), read L50 (i.e., number of contigs whose cumulative length covers half the assembled genome), and the count of benchmark universal single-copy ortholog genes (BUSCOs) present in the assembly. Genome contiguity was visualized using the QUAST 5.3.0 package (https://quast.sourceforge.net).

### Mitochondrial assembly

The mitogenome was extracted and characterized from the reads using MitoHiFi v3.2.2 (https://github.com/marcelauliano/MitoHiFi). MitoHiFi identifies putative mitogenomic contigs by comparing them to reference mitogenomes from related species (Uliano-Silva et al., 2023). We manually specified the reference *Dermacentor silvarum* mitogenome for comparison (NC_026552.1). Default parameters were used with specifications for the invertebrate mitochondrial code. MitoHiFi identified, circularized, and annotated a single putative contig.

### Mitochondrial phylogeny

Our *D. variabilis* mitogenome was compared against others available in NCBI along with related species to build a phylogeny using MAFFT v7.5 for alignment (Katoh et al., 2019). Trees were generated and bootstrapped using Iqtree2 v2.3.6 (https://github.com/iqtree/iqtree2) and visualized using FigTree v1.4.4 (https://github.com/rambaut/figtree/releases).

### Repeats

Initial repeat content identification was performed using RepeatMasker v4.1.4 (https://www.repeatmasker.org/) with the complete Dfam library v3.6 (https://www.dfam.org/home) as described by Flynn et al. (2020) and Storer et al. (2021). RepeatMasker masked a low proportion of the genome, indicating that much of the genome’s repetitive content is not annotated in the public database. Thus, we compiled a custom database of repetitive element content using RepeatModeler2 (https://github.com/Dfam-consortium/RepeatModeler). We then re-ran RepeatMasker with the resulting custom database (via RepeatModeler2) to identify the complete repetitive element content in our assembly (Flynn et al., 2020).

### Methylation

Nanopore sequencing enables the direct detection of base modifications, such as 5-methylcytosine (5mC) and 5-hydroxymethylcytosine (5hmC), by measuring ionic current changes as nucleotides pass through nanopores on a flow cell. These current changes, determined by the molecular weights of modified versus unmodified bases, allow for methylation detection without requiring additional molecular preparation.

The global methylation and hydroxymethylation (5mC and 5hmC) at CpG dinucleotides were determined using the modified base information stored in the basecalled BAM files. Modified BAM files were concatenated and mapped back to our draft assembly using Minimap2 (https://github.com/lh3/minimap2). The modified mapped BAM was converted to bedMethyl format using modkit v0.3.4 (https://github.com/nanoporetech/modkit), and global 5mC and 5hmC percentages were summarized using AWK (https://www.gnu.org/software/gawk/manual/gawk.html#Manual-History).

### Gene annotation

Gene Model Mapper (GeMoMa) v1.9 (Keilwagen et al., 2019) was used for homology-based gene prediction using transcripts from *Ixodes scapularis* (ASM1692078v2) as a reference. BUSCO was executed in protein mode with the arachnida_odb10 database to assess prediction accuracy and completeness.

### Symbiont detection

The total reads were aligned in succession to reference genomes from *Francisella* species (GCF_001275365.1, GCF_003347135.1, GCF_000833355.1) and *Rickettsia rickettsii* (GCF_000018225.1). Breadth and depth were calculated with samtools. Data was visualized with BAMstats v1.25 (https://bamstats.sourceforge.net).

### Data Availability

Detailed computational methods are available in Supplementary file S1. The assembly is available via the NCBI Genome archive at PRJNA1224226, BioSample SAMN46853854; the raw reads are available in the Sequence Read Archive under the same accession.

## Results and Discussion

### Assembly

We generated a total of 83.9 Gb of long read sequence data for the American dog tick (*Dermacentor variabilis*) with an N50 of 3.6 kb and a mean quality of Q19.1. The relatively short N50 was driven by the difficulty in extracting high molecular weight DNA from adult ticks. Our initial Flye run yielded a draft assembly of 2.5 Gb organized into 109,188 contigs with an N50 of 118 kb (Table 1). The initial assembly was cleaned of contaminant sequences (e.g., vector, adapter, and symbiont), purged of duplicate contigs, and gaps were filled to increase contiguity. Our final assembly was 2.15 Gb consisting of 21,695 contigs with an N50 of 445 kb.

**Table 1.** Genome contiguity and quality statistics for each iterative draft assembly for the reported *Dermacentor variabilis* genome.

| Assembly | Size (bp) | Contigs | N50 (bp) | L50 | BUSCO*<br>complete | BUSCO<br>single | BUSCO<br>duplicate | Gaps (n) |
| --- | --- | --- | --- | --- | --- | --- | --- | --- |
| Initial assembly (flye) | 2,509,290,473 | 109,188 | 118,641 | 4,871 | 92.26% | 90.56% | 1.70% | 0 |
| Purged (purge_dups) | 2,192,205,988 | 56,222 | 146,707 | 3,707 | 95.20% | 93.97% | 1.23% | 0 |
| ntLinked | 2,177,726,303 | 27,881 | 438,836 | 1,304 | 95.20% | 93.97% | 1.23% | 89 |
| purged (nogaps) | 2,177,300,621 | 27,970 | 435,334 | 1,318 | 95.20% | 93.97% | 1.23% | 0 |
| decontaminated | 2,148,583,136 | 21,695 | 445,025 | 1,285 | 95.20% | 93.97% | 1.23% | 0 |

\*BUSCOs calculated with the compleasm tool.

Our final assembled genome had a BUSCO completeness score of 95.2%, comparable to three other published *Dermacentor* genomes (Table 2). BUSCO completeness scores can be deconstructed into single- and duplicate-copy percentages that provide more nuanced information about genome assembly quality. Single-copy genes suggest high-quality assembly, while duplicate-copy genes may indicate assembly errors. The other *Dermacentor* genomes had BUSCO scores over 97% with 2-4% duplicate-copy genes, while ours had a lower completeness score but fewer duplicates (<1% after purging). This is notably lower than the >2% duplicates in *D. albipictus* and *D. andersoni* and up to 4% in *D. silvarum*, suggesting potential uncollapsed haplotigs in those assemblies.

**Table 2.** BUSCO scores for the reported *Dermacentor variabilis* genome assembly and the other publicly available *Dermacentor* genome assemblies.

| **Species** | **Build** | **BUSCO\***<br>**complete** | **BUSCO**<br>**single** | **BUSCO**<br>**duplicate** |
| --- | --- | --- | --- | --- |
| Dermacentor variabilis | This report | 95.20% | 93.97% | 1.23% |
| Dermacentor albipictus | USDA_Dalb1.pri | 97.92% | 95.60% | 2.32% |
| Dermacentor silvarum | BIME_Dsil_1.4 | 97.41% | 93.39% | 4.02% |
| Dermacentor andersoni | qqDerAnde1.2 | 97.65% | 95.57% | 2.08% |

Tick genomes are challenging to assemble due to their large size, complexity, and high proportion of repetitive elements. Short-read sequencing technologies often fail to resolve long repetitive sequences, limiting the generation of contiguous tick genomes. Long-read sequencing technologies, such as ONT and PacBio, address these limitations by spanning extensive repetitive regions. Advancements in third-generation sequencing technologies over the past decade have led to a surge in assembled tick genomes, and our assembly contributes to this expansion which significantly enhances our understanding of tick biology.

### Mitochondrial structure and phylogeny

The mitochondrial genome was extracted from the assembly and characterized with MitoHiFi, a tool explicitly designed for identifying and annotating mitochondrial contigs from long-read sequencing data (Uliano-Silva et al., 2023). The mitogenome assembly is a single circular contig spanning 14,845 bp, 6-8 bp longer than reference mitogenomes from various *D. variabilis* isolates on NCBI. The additional base pairs present in our mitogenome may stem from either homopolymer errors, a known limitation of nanopore sequencing, or true indels in our organism. Our mitogenome contained 22 tRNA genes, 13 protein-coding genes, and 2 rRNA genes consistent with the ancestral hard tick configuration (Wang et al., 2019), clustering monophyletically with other *D. variabilis* collected from the Upper Midwest (Figure 1a). Although our mitogenome clustered with other isolates from the same geographical region, all *D. variabilis* mitogenomes were highly similar (>98.9%) over their entire length. Congeneric phylogenetic comparison supported previous analyses, clustering our mitogenome with the *D. variabilis* reference mitogenome (NC_061217), nested within a clade of *Dermacentor spp*. from the Americas, and distinct from closely related European and Asian *Dermacentor spp*. (Reynolds et al., 2023; Figure 1b).

**Figure 1.**
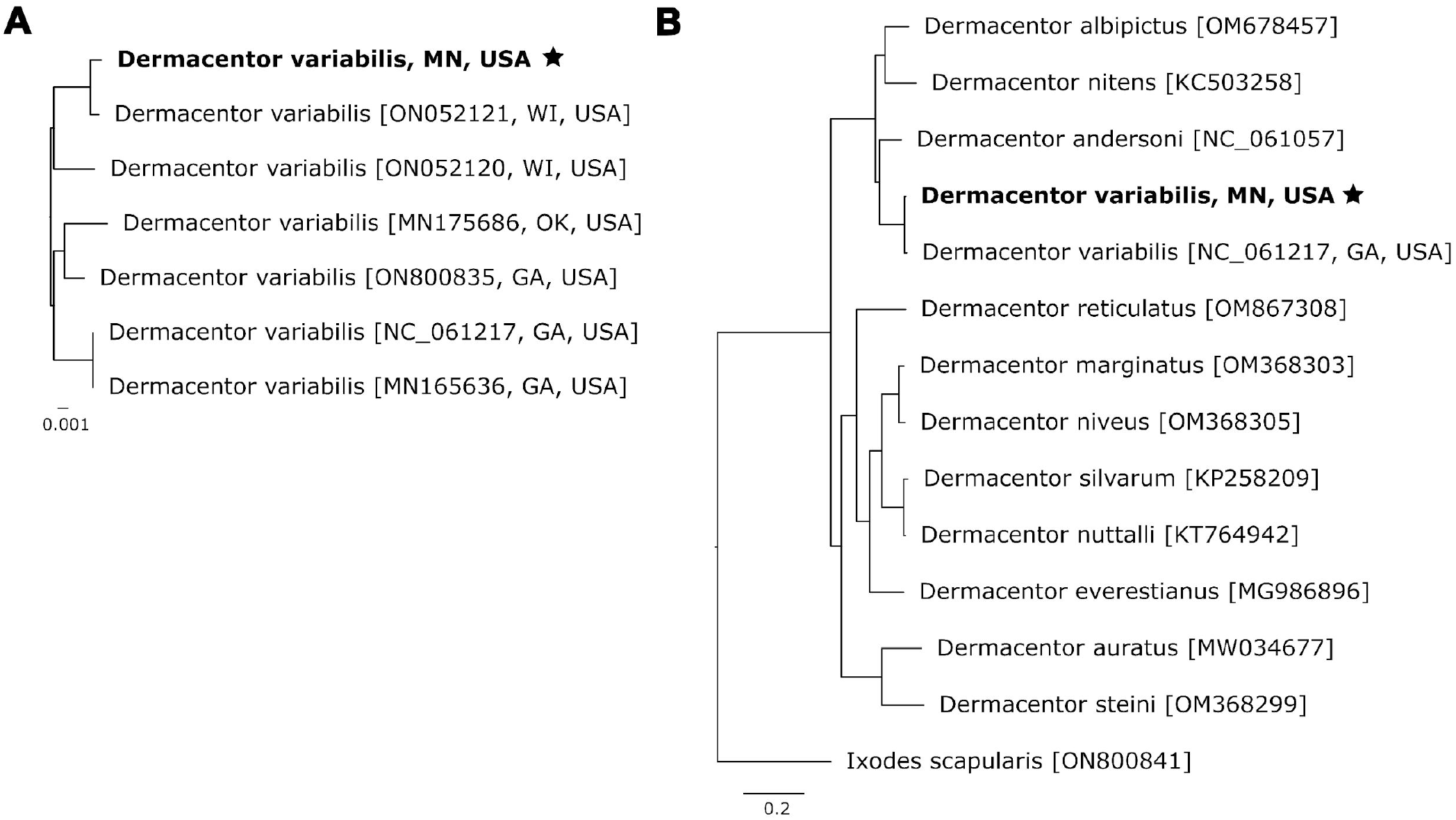
Phylogenetic relationships between *Dermacentor* species were reconstructed using bootstrapped maximum likelihood comparisons of our assembled mitogenome to existing hard tick reference mitogenomes. Phylogenies were created using the entire mitochondrial genomes. (A) depicts relatedness among *D. variabilis* mitogenomes from throughout the United States, and (B) displays relationships between all available *Dermacentor* mitogenomes, rooted with *I. scapularis* as the outgroup.

### Repetitive DNA

Tick repeats are not well characterized in the public Dfam database, limiting our understanding of repetitive element content in the Ixodidae. To address this gap, we leveraged RepeatModeler2 to curate a custom database of *de novo* repeats from our draft assembly. We subsequently used RepeatMasker to mask these repeats in our final assembly. RepeatModeler discovered 4,372 distinct repeat families, 58% categorized as unknown. This finding is consistent with previous reports, which also found that more than 45% of repeats in the *I. scapularis* genome were unclassified (De et al., 2023). When we applied RepeatMasker to our assembly, it classified 61% of the genome as repetitive, with 32% of this content consisting of retroelements, predominately long interspersed nuclear elements (LINEs) and long terminal repeat (LTRs) elements (Table 3). Only 2.5% of repetitive content was categorized as DNA transposons, in stark contrast to other hematophagous insects, such as tsetse flies, where DNA transposons comprise a substantial fraction of repetitive DNA (Attardo et al., 2019). Further, a considerable portion of the repetitive content (24%) remained unclassified, underscoring the complexity of transposable element (TE) dynamics in organisms with genomes dominated by selfish genetic elements. These unclassified repeats highlight the need for further investigation into the nature and role of TEs in tick genomes, which could provide insights into their evolution and genomic plasticity.

**Table 3.**
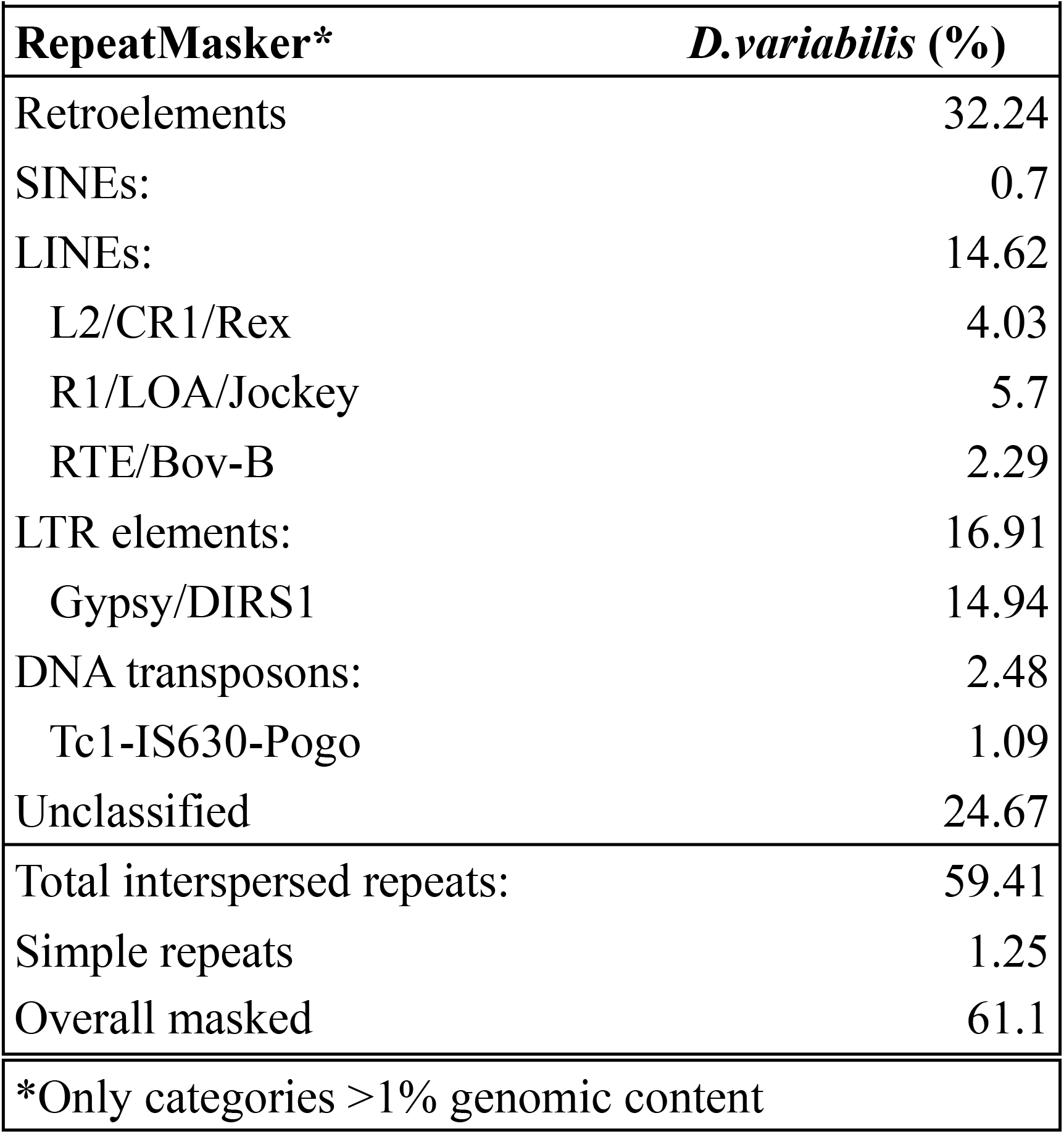
Repetitive DNA content for the reported *Dermacentor variabilis* genome assembly.

|  |  |
| --- | --- |
| **RepeatMasker\*** | ***D.variabilis* (%)** |
| Retroelements | 32.24 |
| SINEs: | 0.7 |
| LINEs: | 14.62 |
| L2/CR1/Rex | 4.03 |
| R1/LOA/Jockey | 5.7 |
| RTE/Bov-B | 2.29 |
| LTR elements: | 16.91 |
| Gypsy/DIRS1 | 14.94 |
| DNA transposons: | 2.48 |
| Tc1-IS630-Pogo | 1.09 |
| Unclassified | 24.67 |
| Total interspersed repeats: | 59.41 |
| Simple repeats | 1.25 |
| Overall masked | 61.1 |
| *Only categories >1% genomic content |  |

We applied our custom database of *D. variabilis* repeat families to mask three other published *Dermacentor* genomes. Our assembly shared a similar repeat content level with *D. albipictus* (59.64%), while *D. andersoni* (62.02%) and *D. silvarum* (54.27%) exhibited higher and lower levels, respectively. The lower repetitive content in our estimate of the *D. silvarum* genome compared to Jia et al. (2020) likely results from differences in the repetitive element databases used (Repbase vs. RepeatModeler).

Repeat content in the *D. variabilis* genome aligns with other metastriate tick genomes, such as *Rhipicephalus sanguineus* (61.6%) and *Haemaphysalis longicornus* (59.3%), but differs from prostriate ticks like *I. scapularis* (69%) and *I. ricinus* (73%), despite belonging to the same family (De et al., 2023; Ronai et al., 2024). These findings reinforce the phylogenetic relationships displayed in Figures 1a and 1b. Lineage-specific repetitive element content, driven by variations in TE activity, genome defense mechanisms (e.g., piRNA clusters), and evolutionary pressures, is evident across arthropod taxa (Petersen et al., 2019). Considering TEs comprise the majority of tick genomes, understanding the forces shaping TE dynamics in hard ticks remains a priority. Detailed information is available in Supplementary file S2.

### Gene annotation

Putative gene predictions were completed using the GeMoMa annotation tool. GeMoMa predicts gene models utilizing existing gene models in related reference species (Keilwagen et al., 2019). We used the *I. scapularis* annotation (ASM1692078v2) as a reference, as it remains the most robust gene model among hard ticks. We detected 21,722 putative genes in our genome assembly. The protein BUSCO score was 80.88% complete (71.20% single copy and 9.68% duplicates), indicating a high level of completeness and reliability in the annotated genome. However, some regions may still contain fragmented or missing orthologs. Our genome annotation would benefit from detailed RNA-seq data and future methodological refinement. Annotations are available in Supplementary file S3.

### DNA Methylation

We observed a genome-wide DNA methylation level of 9.9% in *D. variabilis*. Previous studies on other metastriate ticks, such as *D. silvarum* and *H. longicornus*, have characterized DNA methyltransferases, i.e., enzymes responsible for adding and maintaining methyl marks. Still, this study represents the first report of genome-wide methylation in metastriate ticks (Agwunobi et al., 2021).

The CpG dinucleotide composition of the *D. variabilis* genome suggests balanced transition rates in contrast to the biased CpG depletion observed in the mouse genome (mm39; Figure 2). Mammalian genomes, with global methylation levels exceeding 70%, exhibit accelerated loss of CpG dinucleotides— a pattern not evident in tick genomes. In arthropods like ticks, methylation may contribute to phenotypic variation, particularly in contexts with limited genotypic variation. Recent studies highlight the role of epigenetic mechanisms in hard tick cold tolerance, reproduction, fecundity, and overall fitness (De et al., 2023; Nwanade et al., 2022; Yu et al., 2024). Future research should focus on elucidating the genomic distribution, spatial organization, and functional significance of methylation in ticks.

**Figure 2.**
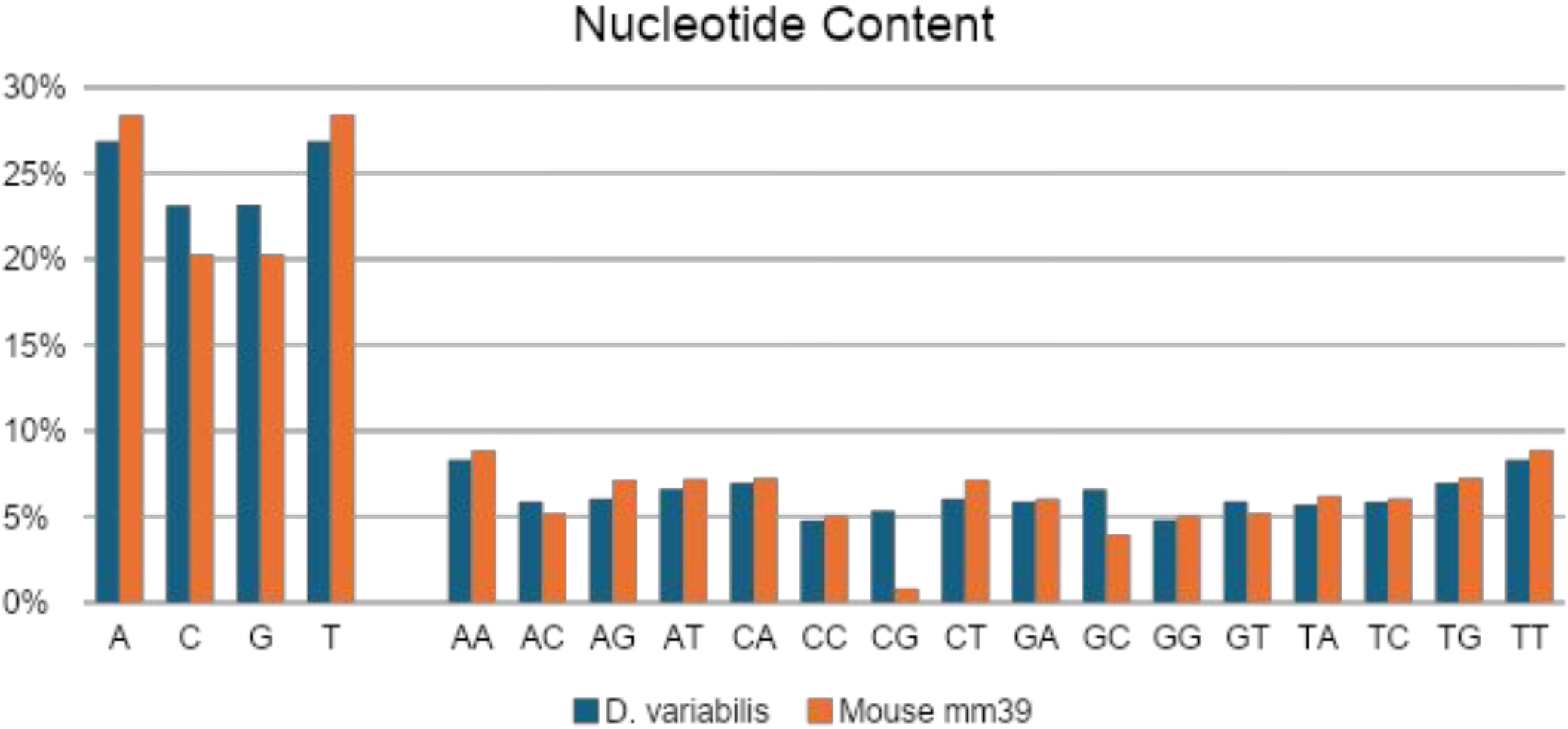
Distribution of nucleotide content comparing *D. variabilis* and *Mus musculus*, the house mouse.

### Symbiont identification

The American dog tick is a potential vector for Tularemia (*Francisella tularensis*) and Rocky Mountain Spotted Fever (*Rickettsia rickettsii*). All reads were aligned to potential reference genomes to confirm the presence of any symbionts, and breadth and depth were calculated. Contig BLAST hits matched to *F. persica, F. opportunistica*, and *F. tularensis*, which were chosen for alignment. The depth and breadth of mapped reads are displayed in Table 4 and Figure 3. Mapped reads demonstrated the highest breadth and depth of mapping to *F. persica*, indicating the presence of only this non-pathogenic endosymbiont.

**Table 4.** Read alignment statistics for pathogenic and non-pathogenic microorganism reads extracted from the *Dermacentor variabilis* raw sequence data.

| **Target** | **Contigs** | **Avg. depth (X)** | **Reference (bp)** | **Length >3X (bp)** | **Breadth (>3X)** | **Present** |
| --- | --- | --- | --- | --- | --- | --- |
| *Francisella persica* | 16 | 20.46 | 1,516,676 | 1,457,338 | 96.09% | Yes |
| *Francisella opportunistica* | 31 | 18.78 | 1,839,233 | 1,443,153 | 78.46% | No |
| *Francisella tularensis* | 10 | 18.47 | 1,870,206 | 1,503,758 | 80.41% | No |
| *Rickettsia rickettsii* | 0 | 6.5 | 1,257,710 | 1,216,534 | 96.73% | Yes |

**Figure 3:**
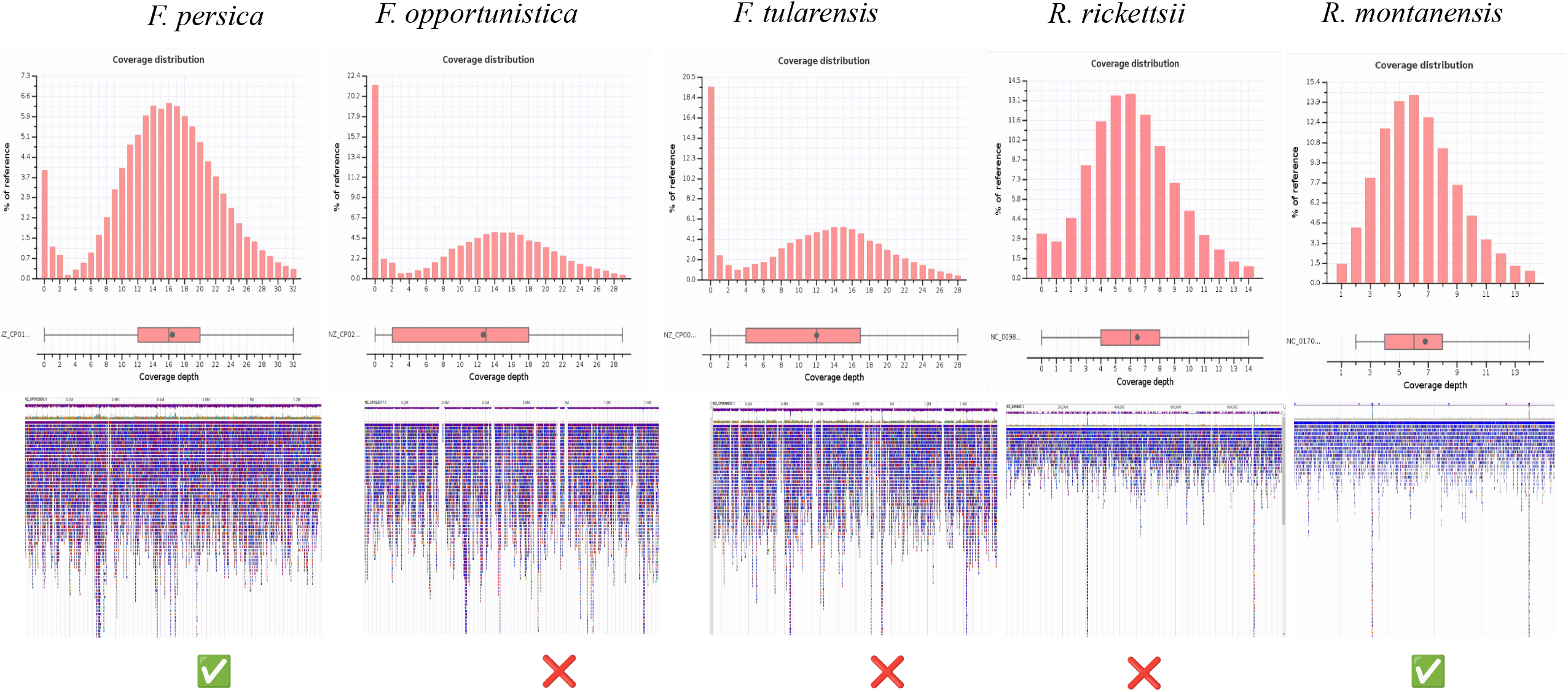
Alignment of mapped reads to the pathogenic and non-pathogenic (endosymbiont) reference genomes. Histograms display the coverage distribution for mapped reads, boxplots display the depth of coverage, and the genome viewer displays mapped read distribution across the reference genomes.

Although no contigs were initially identified as *Rickettsia rickettsii*, we attempted to align reads to *Rickettsia montanensis* to determine its presence or absence, as it has been detected in *D. variabilis* ticks from Minnesota (Hecht et al., 2019). We identified reads mapping to 99% of the reference genome but at a lower depth than the *Francisella*-like endosymbiont, suggesting comparatively low levels of *Rickettsia* in *D. variabilis*. Intriguingly, we could not assemble contigs for this species, preventing definitive identification when using the kraken2 standard library. Difficulty in re-constructing the reads mapping to this genome may have arisen from its inherent lower cell count, inevitably leading to lower genome coverage. This affirms previous reports of *R. montanensis* presence in Minnesota and should be monitored, considering its association with clinical and subclinical symptoms in humans (Vincent and Hulstrand, 2022).

## Conclusions

Our study presents the first whole-genome assembly for *Dermacentor variabilis*, the American dog tick, a hard tick widely distributed across North America. Our assembly achieved a BUSCO completeness score comparable to other publicly available *Dermacentor* genomes, underlining the utility of nanopore sequencing for generating high-quality tick genomes. Beyond its value for understanding tick biology and evolution, this assembly facilitates improved detection of tick-associated microbes, including endosymbionts and potential pathogens.

Amplification-free unbiased detection and identification of tick endosymbiont species within ticks remains challenging, owing to their complex evolutionary histories and low copy number compared to mitochondrial DNA. Nanopore sequencing allowed us to distinguish a *Francisella*-like endosymbiont, mapping more closely to *F. persica* than *F. tularensis*, using read mapping statistics (e.g., breadth and depth) to avoid false positives commonly encountered with PCR-based identification (Kugeler et al., 2005). Moreover, we detected *Rickettsia montanensis* through total read alignment to its reference genome despite an inability to assemble contigs for this species. These findings underscore the need to refine bioinformatic tools for long-read sequence data to better distinguish pathogenic from non-pathogenic bacterial DNA in ticks and highlights the importance of broadening genomic resources to improve our understanding of tick-microbe molecular interactions.

Expanding genomic resources for medically relevant arthropods is critical for uncovering evolutionary processes underlying vectorial capacity, migration, and life-history traits. Ticks, second only to mosquitoes in global vector significance, represent an expanding area for genomic exploration.

Sequencing of hard tick genomes has provided key insights into genome biology, permitting comparative analysis between hard tick groups (i.e., Prostriata and Metastriata) that can reveal lineage-specific evolution of gene families, hematophagy, and ecological adaptations (Cerqueira De Araujo et al., 2025; De et al., 2023; Jia et al., 2020). Despite over 450 documented metastriate species, only ten unique genome assemblies are available on NCBI. Our *D. variabilis* genome helps close this gap, offering a critical resource for comparative analyses of tick genetic diversity, evolutionary history, and population genetics as ticks expand into new geographic areas and pose emerging zoonotic threats.

## Author contributions

This work was a collaborative effort by the members of the graduate course ANSC 8520 taught by Dr. Faulk in the Department of Animal Science at the University of Minnesota in the Fall of 2024. Students were Matt Villalta, Saul Aguirre, Lauren Ecklund, Trek Stenger, Idil Abdi, and Sree Venigalla. Students performed analyses and provided text for the manuscript. Jacob Cassens provided samples, performed analyses, and wrote the manuscript. Ms. Shiffman provided additional samples. Drs. Kristen Bastug and Beth Thielen completed assemblies and analyses and edited the manuscript. Dr. Faulk performed analyses and wrote and edited the manuscript. All authors have read and approved the manuscript prior to submission.

## Data Availability Statement

Supplementary File S1 provides detailed descriptions of the computational steps outlined in the methods. Supplementary File S2 contains the transposon family database generated from RepeatModeler2.

Supplementary File S3 contains the gene annotations in GFF format. Sequence data for the whole genome and mitogenome assemblies are deposited in GenBank with the accession number JBLSAA000000000.

## Acknowledgments

The authors thank William Finical for kindly providing the ticks.

## Supplementary Files

**File S1:** Supplementary methods

**File S2:** Transposon family database

**File S3:** Annotations in GFF format

